# DNA damage response signaling to mitochondria drives senescence

**DOI:** 10.1101/2022.09.22.509001

**Authors:** Shota Yamauchi, Yuki Sugiura, Junji Yamaguchi, Xiangyu Zhou, Takeru Odawara, Shunsuke Fukaya, Isao Naguro, Yasuo Uchiyama, Hidenori Ichijo

## Abstract

Cellular senescence is a stress-induced irreversible cell cycle arrest typically accompanied by expression of the cyclin-dependent kinase inhibitor p16^INK4a^ (hereafter referred to as p16) and mitochondrial dysfunction^1^. Recent studies have indicated that p16-expressing senescent cells accumulate in the body over time and contribute to aging^1, 2^. Many stresses, such as telomere shortening and oncogene activation, induce senescence by damaging nuclear DNA^1^. However, the molecular mechanisms linking DNA damage to senescence remain unclear. Here, we show that the outer mitochondrial transmembrane protein BNIP3 drives senescence by triggering a DNA damage response (DDR) of mitochondria. BNIP3 was identified in a genome-wide siRNA screen for genes required for p16 expression upon DNA damage. Mass spectrometric analysis of BNIP3-interacting proteins yielded the DDR kinase ATM and subunits of the mitochondrial contact site and cristae organizing system (MICOS) complex. BNIP3 is an ATM substrate that increases the number of mitochondrial cristae upon DNA damage. This increase enhances the oxidation of fatty acids to acetyl-CoA, an acetyl group donor, thereby promoting histone acetylation and associated p16 expression. Our findings indicate that DDR signaling to mitochondria promotes p16 expression by altering mitochondrial structure and metabolism and highlight the importance of nuclear–mitochondrial communication in senescence induction.

---

Cellular senescence is a tumor-suppressive mechanism that prevents the proliferation of potentially tumorigenic cells^1, 3^. However, recent studies have indicated that it also contributes to aging and age-related diseases^1^. The expression of p16, a marker and effector of senescence, increases in various tissues with age^4, 5^. Increased p16 expression decreases the tissue-regenerative potential of pancreatic β-cells and neuronal progenitor cells^6, 7^. p16-expressing cells that accumulate in the body impair heart and kidney function and shorten the healthy lifespan^2^. Furthermore, the *INK4/ARF* locus, encoding p16, is a hotspot for susceptibility single nucleotide polymorphisms associated with age-related diseases, such as atherosclerosis and diabetes^1^.

Cellular senescence is a DDR^1^. Many senescence-inducing stresses produce DNA damage, mainly double-strand breaks, and activate the DDR kinase ATM. ATM stabilizes and activates the transcription factor p53, which promotes expression of the cyclin-dependent kinase inhibitor p21. p21 arrests the cell cycle transiently to provide time for DNA repair^8^. If DNA damage persists, the resulting prolonged DDR signaling promotes the expression of p16^1, 8^. Numerous transcription factors and histone-modifying enzymes have been reported to contribute to p16 expression^9^. However, little is known about the molecular mechanisms linking DNA damage to p16 expression.

## BNIP3 links DNA damage to p16 expression

Treatment with doxorubicin, which produces DNA double-strand breaks, increased p16 expression in IMR-90 normal human fibroblasts (Extended Data Fig. 1a–d). This increase was suppressed by ATM knockdown but was augmented by p53 knockdown^4, 10^, suggesting that downstream effectors of ATM other than p53 mediate DNA damage-induced p16 expression. By combining anti-p16 immunofluorescence and automated microscopy, we established a p16 quantification assay that can be used in large-scale screens^11^ (Extended Data Fig. 1e). The specificity of the p16 antibody was confirmed with a p16 siRNA (Extended Data Fig. 1f). We used this assay to carry out a genome-wide siRNA screen for genes required for p16 expression upon DNA damage (Fig. 1a). IMR-90 cells were transfected with siRNAs in 384-well plates. p16 expression was induced by doxorubicin treatment and quantified on Day 10. Genes whose knockdown significantly suppressed p16 expression included *BNIP3*, which encodes an outer mitochondrial transmembrane protein previously implicated in cell death and mitophagy^12^, and *NDUFA8*, which encodes an essential subunit of respiratory chain complex I^13^ (Fig. 1b).

**Fig. 1.**
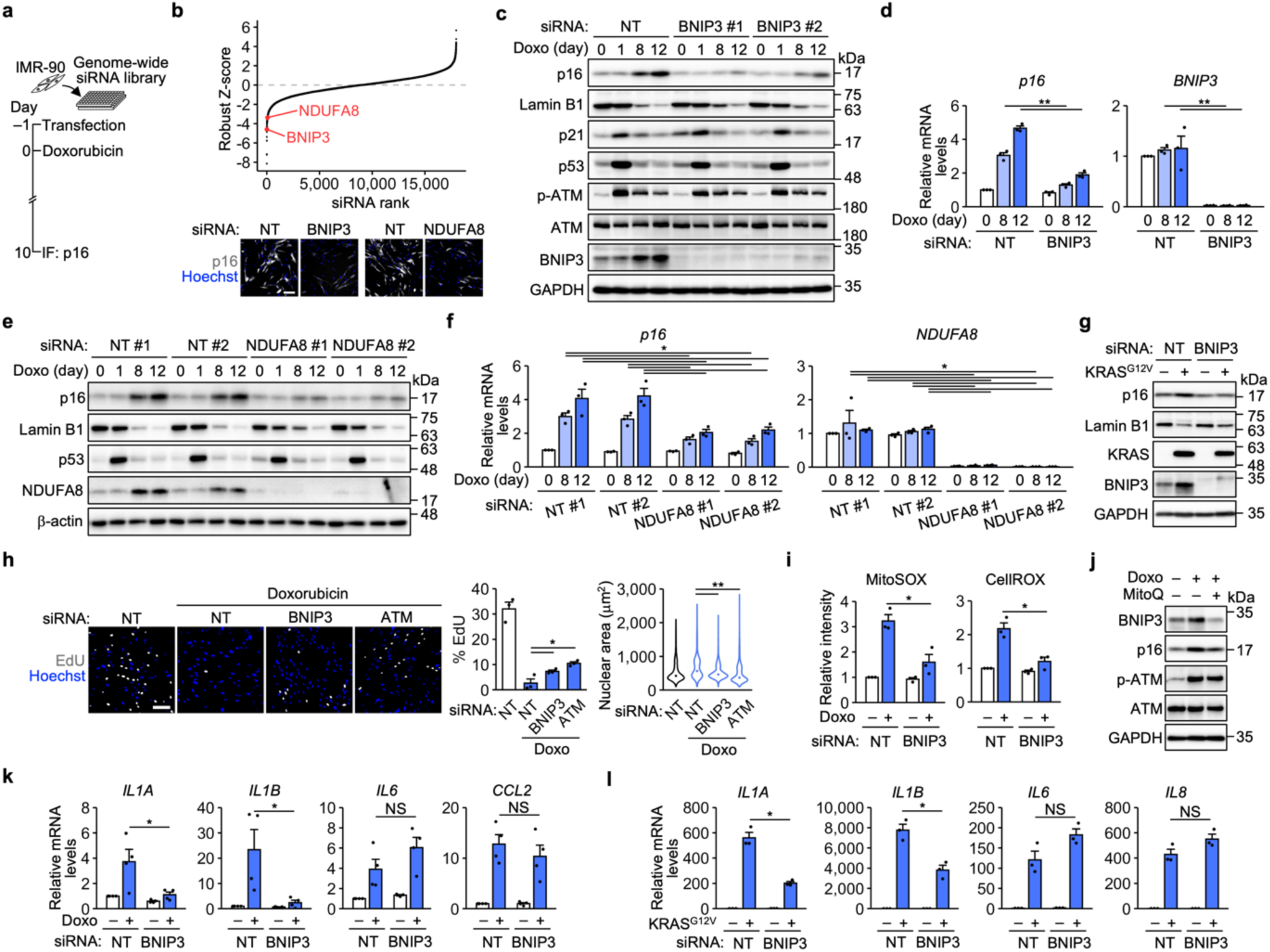
A genome-wide siRNA screen implicates BNIP3 in DNA damage-induced senescence. **a**, Schematic of the siRNA screen. IF, immunofluorescence. **b**, Top, robust Z-scores of the screened siRNAs. Bottom, images of IMR-90 cells transfected with nontargeting (NT), BNIP3, and NDUFA8 siRNAs from the siRNA screen. Scale bar, 200 μm. **c**–**f**, Immunoblot (**c**, **e**) and qPCR (**d**, **f**) analyses of IMR-90 cells transfected with the indicated siRNAs and treated with doxorubicin (Doxo). Representative of three independent experiments (**c**, **e**). *n* = 3 independent experiments (**d**, **f**). **g**, Immunoblot analysis of IMR-90 cells expressing ER-KRAS^G12V^ for 10 days. Representative of three independent experiments. **h**, Left, EdU assay using IMR-90 cells treated with 50 ng ml^−1^ Doxo for 24 h and incubated in fresh medium for 7 days. Scale bar, 200 μm. Center, percentage of EdU-positive cells. *n* = 3 biological replicates. Right, distribution of the nuclear area. The dots represent the median values. *n* = 3215 (siNT), 5487 (siNT + Doxo), 4802 (siBNIP3 + Doxo), 7857 (siATM + Doxo) cells. **i,** ROS measurements with the indicated probes. *n* = 3 independent experiments. **j**, Immunoblot analysis of IMR-90 cells treated with Doxo and MitoQ for 12 days. Representative of three independent experiments. **k**, qPCR analysis of IMR-90 cells treated with Doxo for 12 days. *n* = 4 independent experiments. **l**, qPCR analysis of IMR-90 cells expressing ER-KRAS^G12V^ for 10 days. *n* = 3 independent experiments. The bars represent the means ± s.e.m. Statistical analysis was performed using unpaired two-tailed Student’s *t*-test (**d**, **i**, **k**, **l**), Tukey’s multiple comparison test (**f**), Dunnett’s multiple comparison test (**h**, center), and the Wilcoxon rank-sum test with Bonferroni correction (**h**, right). \**P* < 0.05, ** *P* < 0.01. NS, not significant.

Mitochondria alter their structure and function during senescence and contribute to senescence features, such as p16 expression, cell enlargement, reactive oxygen species (ROS) accumulation, and cytokine secretion^14–20^. We confirmed by immunoblotting and quantitative PCR (qPCR) that knockdown of BNIP3 or NDUFA8 suppressed doxorubicin-induced p16 expression (Fig. 1c–f). Doxorubicin treatment increased BNIP3 and NDUFA8 levels along with p16 levels (Fig. 1c, e). BNIP3 knockdown only slightly attenuated the loss of lamin B1, a senescence marker^1^, and did not markedly affect activation of the ATM–p53–p21 pathway (Fig. 1c). BNIP3 knockdown also suppressed p16 expression during oncogenic RAS (ER-KRAS^G12V^)-induced senescence (Fig. 1g). In a mouse mammary tumor model, BNIP3 has been found to prevent tumor cell proliferation and tumor growth^21^. We examined whether BNIP3 is involved in senescence-associated proliferation arrest. Doxorubicin treatment decreased the percentage of EdU-positive proliferating cells and concomitantly increased nuclear size (Fig. 1h). These changes were attenuated by BNIP3 or ATM knockdown.

BNIP3 has been implicated in mitochondrial ROS production during cell death^12^. BNIP3 knockdown suppressed an increase in mitochondrial ROS during doxorubicin-induced senescence (Fig. 1i). Conversely, scavenging of mitochondrial ROS with MitoQ, a mitochondria-targeted antioxidant, prevented an increase in BNIP3 (Fig. 1j), suggesting that BNIP3 and mitochondrial ROS form a positive feedback loop. However, the effect of MitoQ on p16 expression was limited. Senescent cells secrete inflammatory cytokines and chemokines^22–24^. This feature is termed the senescence-associated secretory phenotype (SASP) and is a potential therapeutic target in age-related diseases^1^. BNIP3 knockdown selectively suppressed expression of IL-1α and IL-1β during both doxorubicin- and oncogenic RAS-induced senescence (Fig. 1k, l). These results indicate that BNIP3 is involved in p16 expression and several other senescence features.

## ATM phosphorylates BNIP3 in mitochondria

BNIP3 is a transmembrane protein of the outer mitochondrial membrane^12^. We confirmed that BNIP3 was localized to mitochondria in both proliferating and doxorubicin-induced senescent cells (Fig. 2a). The mechanisms by which DNA damage affects mitochondria during senescence remain elusive^14^. p53 can regulate mitochondrial proteins^25^, but we found that it was dispensable for p16 expression^4^ (Extended Data Fig. 1d). To investigate the regulation and function of BNIP3, we identified BNIP3-interacting proteins by mass spectrometry. FLAG-BNIP3 was overexpressed in HEK293T cells and immunoprecipitated with anti-FLAG beads. The coprecipitated proteins included ATM and ATR, a DDR kinase that is activated by single-stranded DNA and shares many substrates with ATM^26^. ATM is localized in part to mitochondria and contributes to the maintenance of respiratory activity^27^, possibly by phosphorylating unidentified mitochondrial substrates^26^. Subcellular fractionation of IMR-90 cells showed the presence of ATM in mitochondria (Fig. 2b). Notably, doxorubicin treatment for 24 h increased phosphorylated ATM levels in the mitochondrial and cytosolic fractions, suggesting that a portion of ATM activated in the nucleus translocates to mitochondria.

**Fig. 2.**
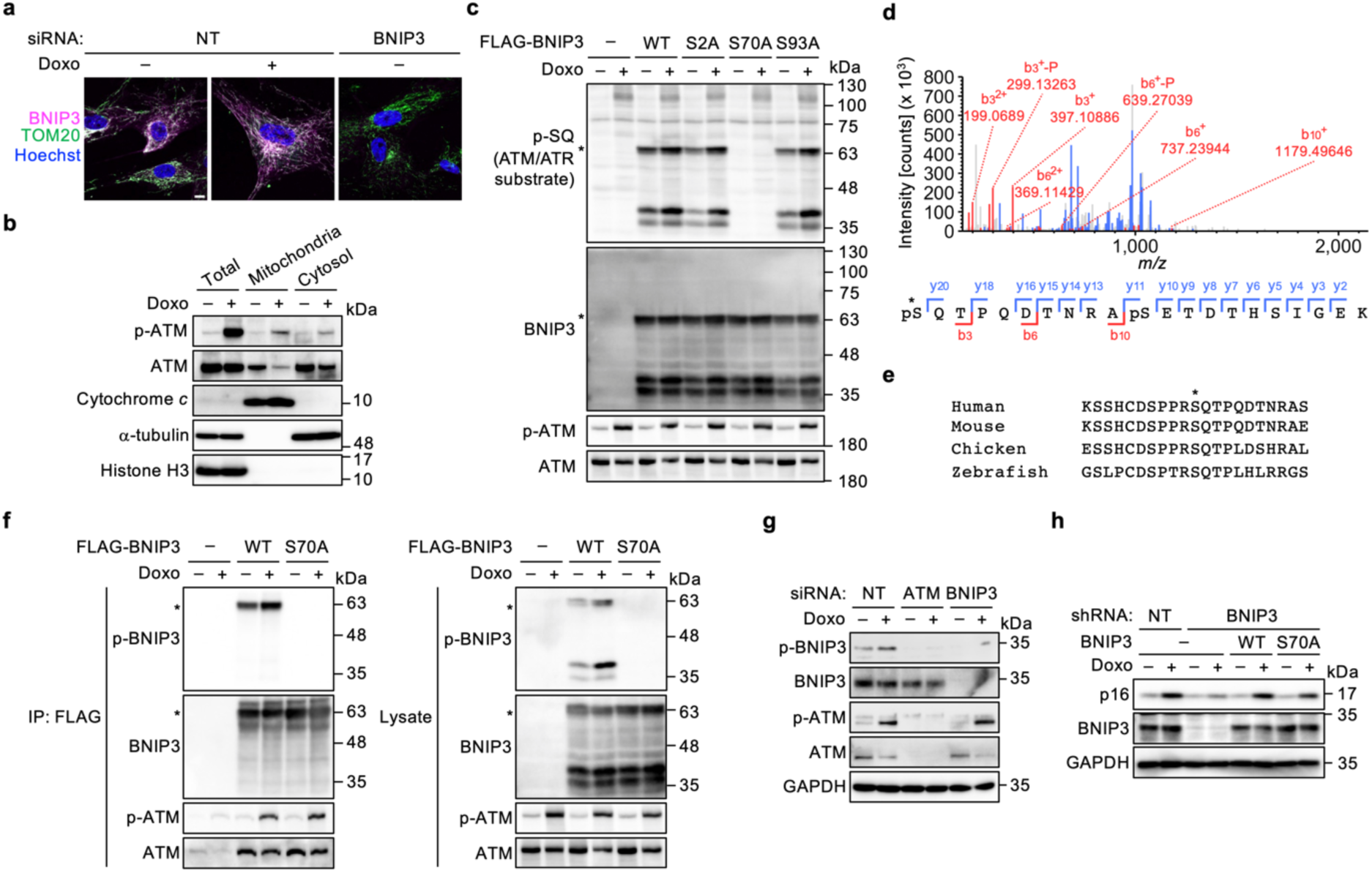
BNIP3 is a mitochondrial substrate of ATM. **a**, Immunofluorescence analysis of IMR-90 cells transfected with nontargeting (NT) and BNIP3 siRNAs and treated with doxorubicin (Doxo) for 12 days. Scale bar, 10 μm. Representative of three independent experiments. **b**, Subcellular fractionation of IMR-90 cells treated with Doxo for 24 h. Representative of three independent experiments. **c**, Immunoblot analysis of HEK293T cells transfected with the indicated plasmids and treated with Doxo for 24 h. The asterisks denote the BNIP3 dimer. Representative of three independent experiments. **d**, Top, product ion spectrum of a BNIP3-derived peptide containing phosphorylated S70. Bottom, identified b- and y-ions. The asterisk denotes S70. Representative of two independent experiments. **e**, Alignment of BNIP3 amino acid sequences from the indicated species. The asterisk denotes S70. **f**, Immunoprecipitation (IP) of FLAG-BNIP3 from HEK293T cells treated with Doxo for 24 h. The asterisks denote the BNIP3 dimer. Representative of three independent experiments. **g**, Immunoblot analysis of IMR-90 cells transfected with the indicated siRNAs and treated with Doxo for 24 h. Representative of two independent experiments. **h**, Immunoblot analysis of IMR-90 cells infected with the indicated lentiviruses and treated with 100 ng ml^−1^ Doxo for 12 days. Representative of two independent experiments.

We examined the possibility that BNIP3 is a mitochondrial substrate of ATM. The substrate motifs of ATM and ATR are both SQ/TQ^26^. A phospho-SQ antibody recognized BNIP3 overexpressed in HEK293T cells (Fig. 2c). Mutational analysis indicated that of the three candidate residues, S70 was the phosphorylation site. Phosphorylation at this site was also detected by mass spectrometry (Fig. 2d). S70 and its surrounding sequences are conserved among species (Fig. 2e). We raised a phospho-specific antibody against S70 of BNIP3. As expected, this antibody recognized immunoprecipitated FLAG-BNIP3 in an S70-dependent manner (Fig. 2f). Coprecipitation of ATM was also confirmed. We then investigated the phosphorylation state of endogenous BNIP3. Treatment with doxorubicin for 24 h increased the phosphorylation of BNIP3 at S70 in IMR-90 cells (Fig. 2g). This increase was suppressed by ATM knockdown. To examine the significance of BNIP3 phosphorylation, BNIP3^WT^ and BNIP3^S70A^ were reexpressed in BNIP3-knockdown cells. Doxorubicin-induced p16 expression was restored almost fully by BNIP3^WT^ but only partially by BNIP3^S70A^ (Fig. 2h). These results suggest that BNIP3 is an ATM substrate that mediates DNA damage-induced p16 expression.

## BNIP3 remodels mitochondrial cristae

BNIP3 has recently been regarded as a mitophagy receptor that interacts with LC3 family proteins^21^. However, mitophagy is impaired during senescence^14^, perhaps through mitochondrial elongation^28, 29^. To explore BNIP3 function, we performed Gene Ontology (GO) analysis of the BNIP3-interacting proteins. This analysis revealed the most significant enrichment for ‘‘inner mitochondrial membrane organization’’ proteins, including the MICOS complex subunits MIC60, MIC19, MIC13, and MIC27 and their interacting proteins TMEM11, SAM50, and DNAJC11 (Fig. 3a). MICOS is an inner mitochondrial membrane complex with dual functions: formation of contact sites with the outer membrane and maintenance of the architecture of inner membrane invaginations known as cristae^30^. Although a recent study identified BNIP3 as a TMEM11-interacting protein^31^, the relationship between BNIP3 and MICOS has not been tested. We confirmed that MIC60 and MIC19 coprecipitated with FLAG-BNIP3 by immunoblotting (Fig. 3b). MIC60 is a core subunit whose loss destabilizes the other subunits and MICOS-interacting proteins^30^. MIC60 knockdown decreased BNIP3 levels along with MIC19 levels in IMR-90 cells (Fig. 3c).

**Fig. 3.**
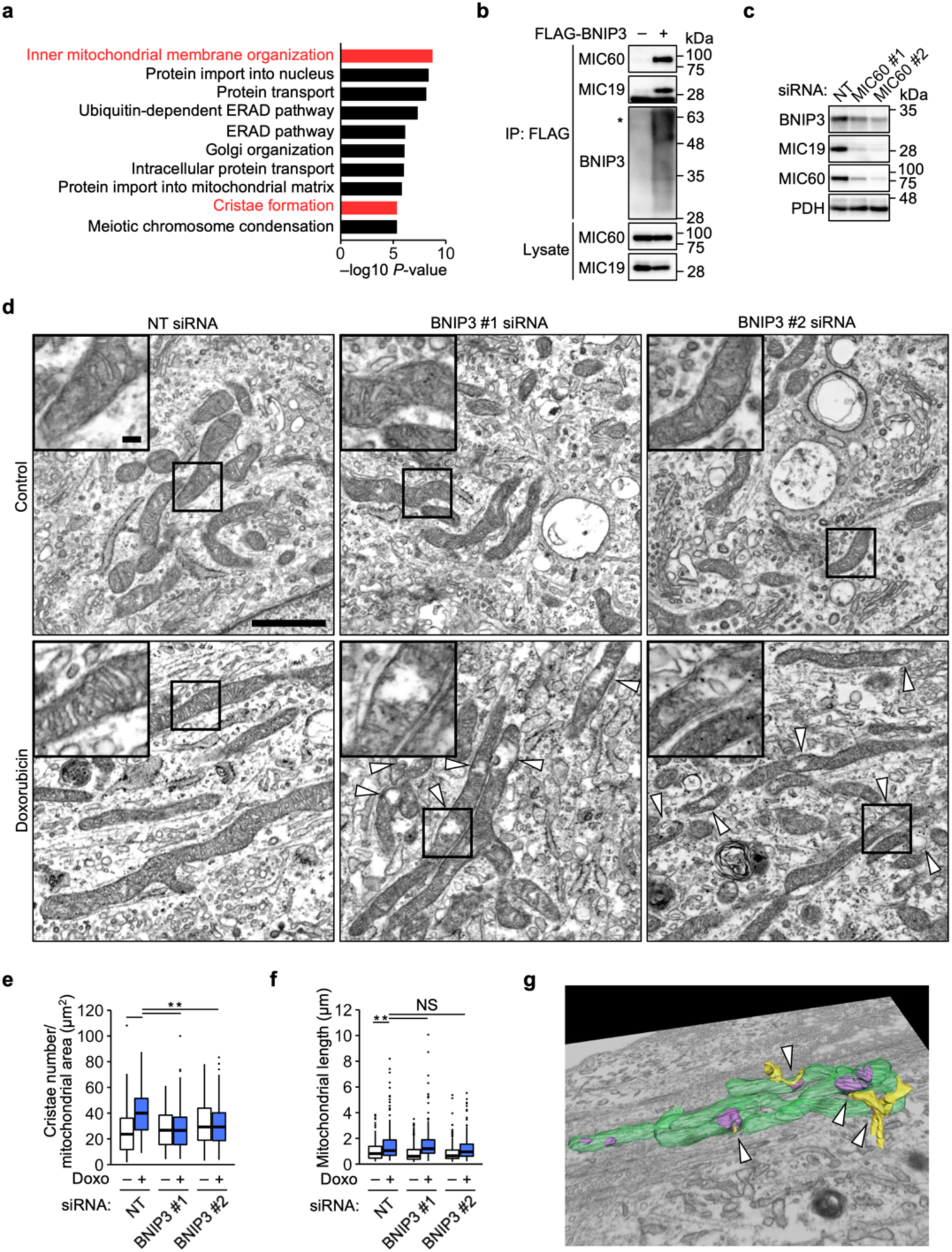
BNIP3 interacts with MICOS and remodels mitochondrial cristae. **a**, GO analysis of BNIP3-interacting proteins identified by mass spectrometry. Representative of two independent experiments. **b**, Immunoprecipitation (IP) of FLAG-BNIP3 from HEK293T cells. The asterisk denotes the BNIP3 dimer. Representative of three independent experiments. **c**, Immunoblot analysis of IMR-90 cells transfected with nontargeting (NT) and MIC60 siRNAs and treated with doxorubicin (Doxo) for 12 days. Representative of three independent experiments. **d**, Electron microscopy of IMR-90 cells transfected with NT and BNIP3 siRNAs and treated with Doxo for 12 days. The arrowheads denote degenerated regions. Scale bar, 1 μm; inset scale bar, 100 nm. Representative of two independent experiments. **e**, **f**, Quantification of the number of cristae per mitochondrial area (**e**) and mitochondrial length (**f**). *n* = 217 (siNT), 149 (siNT + Doxo), 286 (siBNIP3 #1), 173 (siBNIP3 #1 + Doxo), 168 (siBNIP3 #2), 203 (siBNIP3 #2 + Doxo) mitochondria. Center line, median; box limits, upper and lower quartiles; whiskers, 1.5× interquartile range; points, outliers. Statistical analysis was performed using the Wilcoxon rank-sum test with Bonferroni correction. ** *P* < 0.01. NS, not significant. **g**, FIB–SEM of IMR-90 cells transfected with BNIP3 #1 siRNA and treated with Doxo for 12 days. Green, mitochondria; magenta, degenerated regions; yellow, ER. The arrowheads denote mitochondria–ER contact sites. *n* = 1 biological replicate.

Using electron microscopy, we examined whether BNIP3 alters mitochondrial structure during DNA damage-induced senescence. We found that doxorubicin treatment increased the number of cristae per mitochondrial area (Fig. 3d, e). This increase was suppressed by BNIP3 knockdown with partial degeneration of mitochondria. Focused ion beam scanning electron microscopy (FIB–SEM) revealed that the degenerated regions tended to be adjacent to ER–mitochondria contact sites (Fig. 3g), where MICOS has been proposed to facilitate lipid trafficking^30^. Doxorubicin treatment also induced mitochondrial elongation (Fig. 3d, f), as reported in replicative senescence^28^. This elongation was unaffected by BNIP3 knockdown. These results suggest that BNIP3 is a MICOS-interacting protein that remodels mitochondrial cristae during DNA damage-induced senescence.

## BNIP3 alters mitochondrial metabolism

Mitochondrial cristae are the sites of oxidative phosphorylation and house respiratory chain complexes and ATP synthase^30^. Respiratory chain complex I was implicated in p16 expression by our siRNA screen (Fig. 1b, e, f). When generating a proton gradient for ATP synthesis, complex I oxidizes NADH to NAD^+^ (ref. 32). NAD^+^ supply by complex I is essential for mitochondrial metabolic pathways, such as the tricarboxylic acid (TCA) cycle and fatty acid oxidation (FAO)^32, 33^. Mitochondrial metabolites influence gene expression as substrates and cofactors for histone- and DNA-modifying enzymes^32^. We reasoned that BNIP3-dependent cristae remodeling promotes p16 expression by altering mitochondrial metabolism. Metabolomic analysis showed that doxorubicin treatment markedly increased the levels of the TCA cycle intermediates citrate and isocitrate without increasing the levels of pyruvate (Fig. 4a), a glycolytic product that enters the TCA cycle via conversion to acetyl-CoA (Fig. 4b). Doxorubicin treatment also increased the levels of other acetyl-CoA-derived metabolites, such as acetylcarnitine, hexosamine pathway metabolites, and acetylated amino acids (Fig. 4a). These increases tended to be suppressed by knockdown of BNIP3, ATM, or the complex I subunit NDUFA8, raising the possibility that acetyl-CoA metabolism is involved in p16 expression.

**Fig. 4.**
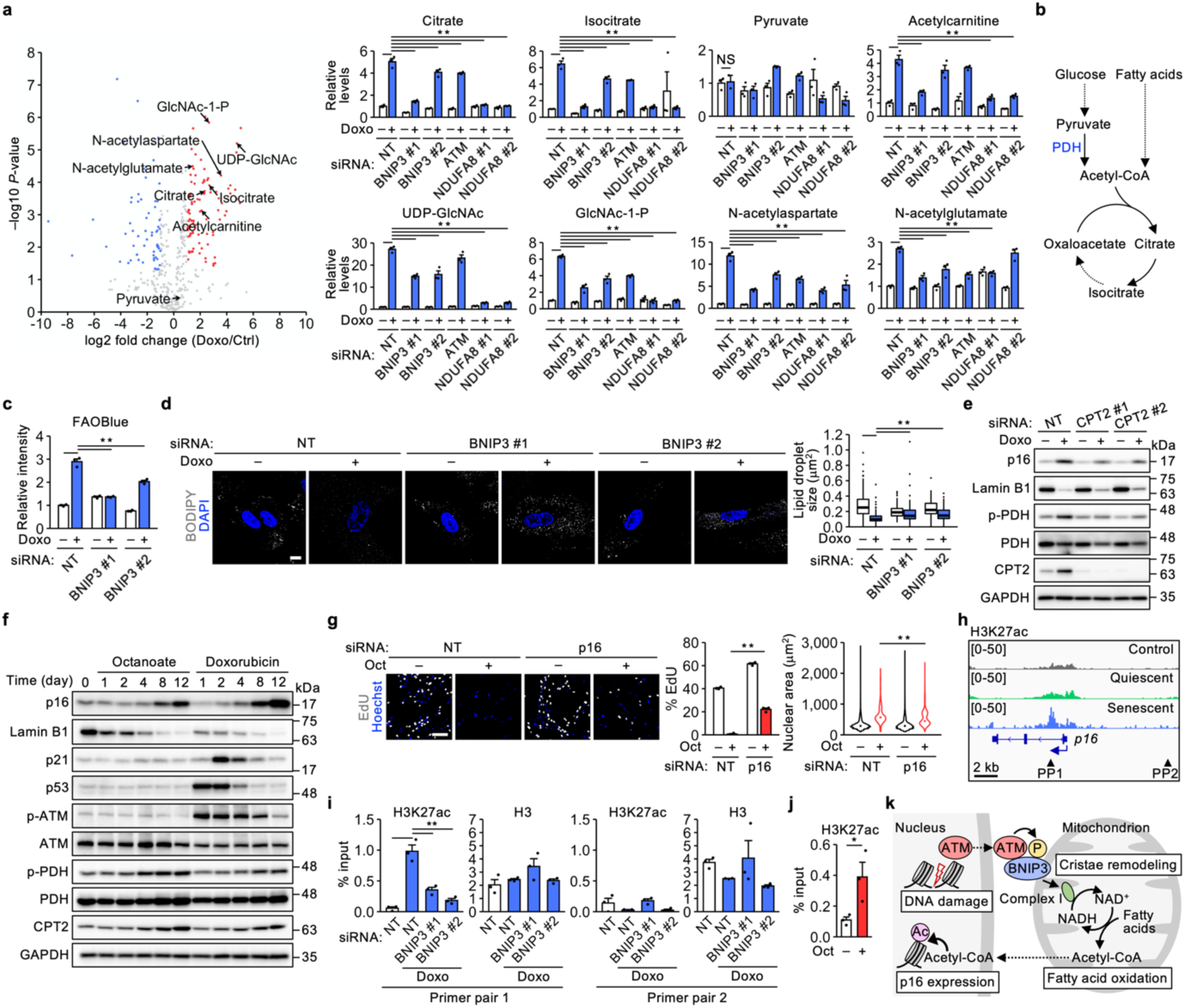
BNIP3 alters mitochondrial metabolism and promotes histone acetylation downstream of the *p16* TSS. **a**, Left, volcano plot of metabolite level changes upon doxorubicin (Doxo) treatment for 12 days in IMR-90 cells transfected with nontargeting (NT) siRNA. Right, relative metabolite levels. *n* = 3 biological replicates. **b**, Schematic of acetyl-CoA and the TCA cycle. **c**, FAO activity assay using IMR-90 cells transfected with NT and BNIP3 siRNAs and treated with Doxo for 12 days. *n* = 3 biological replicates. **d**, Left, lipid droplet staining in IMR-90 cells treated with Doxo for 14 days. Scale bar, 10 μm. Right, quantification of lipid droplet size. *n* = 171 (siNT), 85 (siNT + Doxo), 53 (siBNIP3 #1), 121 (siBNIP3 #1 + Doxo), 102 (siBNIP3 #2), 79 (siBNIP3 #2 + Doxo) cells. Center line, median; box limits, upper and lower quartiles; whiskers, 1.5× interquartile range; points, outliers. **e**, Immunoblot analysis of IMR-90 cells transfected with NT and CPT2 siRNAs and treated with Doxo for 12 days. Representative of three independent experiments. **f**, Immunoblot analysis of IMR-90 cells treated with octanoate (Oct) and Doxo. Representative of three independent experiments. **g**, Left, EdU assay using IMR-90 cells transfected with NT and p16 siRNAs and treated with Oct for 12 days. Scale bar, 200 μm. Center, percentage of EdU-positive cells. *n* = 3 biological replicates. Right, distribution of the nuclear area. The dots represent the median values. *n* = 6766 (siNT), 1005 (siNT + Oct), 9551 (sip16), 2832 (sip16 + Oct) cells. **h**, H3K27ac ChIP–seq analysis of HRAS^G12V^-induced senescent cells. **i,** ChIP‒qPCR analysis of IMR-90 cells treated with Doxo for 12 days using primer pairs (PPs) targeting the regions indicated in **h**. *n* = 3 independent experiments. **j**, ChIP‒qPCR analysis of IMR-90 cells treated with Oct for 14 days using PP1. *n* = 3 independent experiments. **k**, Model for the role of BNIP3 in senescence induction. The bars represent the means ± s.e.m. Statistical analysis was performed using Dunnett’s multiple comparison test (**a**, **c**, **i**), the Wilcoxon rank-sum test with Bonferroni correction (**d**, **g**, right), and unpaired two-tailed Student’s *t*-test (**g**, center, **j**). \**P* < 0.05, ** *P* < 0.01. NS, not significant.

FAO is a complex I-dependent metabolic pathway in which acetyl-CoA is produced from the breakdown of fatty acids^33^. FAO activation during differentiation and starvation involves mitochondrial elongation and an increase in the number of cristae^34–37^. We examined the role of FAO in DNA damage-induced senescence. Doxorubicin treatment increased FAO activity, as reported in oncogenic RAS-induced senescence^38^, and decreased the size of lipid droplets (Fig. 4c, d), storage organelles for fatty acids and sterols^36^. These changes were suppressed by BNIP3 knockdown^39^. Moreover, doxorubicin-induced p16 expression was attenuated by knockdown of CPT2 (Fig. 4e), a mitochondrial enzyme essential for FAO^33^. These results suggest that FAO activation by BNIP3 promotes p16 expression. We noticed that doxorubicin treatment increased CPT2 levels (Fig. 4e). This increase may also contribute to FAO activation.

FAO activation decreases acetyl-CoA production from glucose and sustains fatty acid utilization^40^. This effect, known as the Randle cycle, is primarily due to inhibitory phosphorylation of pyruvate dehydrogenase (PDH), the mitochondrial enzyme that converts pyruvate to acetyl-CoA (Fig. 4b). Doxorubicin treatment increased PDH phosphorylation (Fig. 4e and Extended Data Fig. 2a, b). This increase was attenuated by knockdown of CPT2 or ATM but not by knockdown of p16. In addition, isotope labeling with [U-^13^C]glucose indicated that the conversion of pyruvate to acetyl-CoA and then to citrate and acetylated amino acids decreased during doxorubicin-induced senescence (Extended Data Fig. 2c). These results suggest that PDH is inhibited during DNA damage-induced senescence and are consistent with FAO activation.

To further investigate the importance of FAO in senescence induction, we used the medium-chain fatty acid octanoate. Unlike the more abundant long-chain fatty acids, octanoate freely crosses mitochondrial membranes^41^. When added to cells, octanoate is readily oxidized to acetyl-CoA, an acetyl group donor, and promotes histone acetylation and the associated expression of fatty acid-metabolizing enzymes. Treatment of IMR-90 cells with octanoate increased p16 and phosphorylated PDH levels and decreased lamin B1 levels (Fig. 4f and Extended Data Fig. 3a). In contrast, the ATM–p53–p21 pathway was largely unaffected (Fig. 4f). Octanoate treatment increased CPT2 levels, suggesting that the increase in CPT2 caused by doxorubicin is due to FAO activation (Fig. 4e and Extended Data Fig. 2a, b). Octanoate treatment decreased the percentage of EdU-positive cells and increased nuclear size (Fig. 4g and Extended Data Fig. 3b). These changes were suppressed by p16 knockdown (Fig. 4g). In addition, octanoate treatment increased the percentage of senescence-associated β-galactosidase (SA-β-gal)-positive cells and the expression of inflammatory cytokines and chemokines (Extended Data Fig. 3c, d). These results suggest that FAO activation alone can promote senescence features, including p16 expression.

## BNIP3 promotes histone acetylation

It has long been known that pan-histone deacetylase inhibitors induce senescence-like proliferation arrest by promoting p16 expression^42^. To examine the possibility that FAO activation by BNIP3 promotes p16 expression through histone acetylation, we analyzed public chromatin immunoprecipitation sequencing (ChIP–seq) data. This analysis revealed that acetylation of histone H3 at lysine 27 (H3K27ac) increased downstream of the *p16* transcription start site (TSS) during oncogenic RAS-induced senescence^43^ (Fig. 4h). In general, increased H3K27ac downstream of TSSs is associated with transcription activation^44^. ChIP‒ qPCR analysis indicated that doxorubicin treatment also increased H3K27ac downstream of the *p16* TSS (Fig. 4i). This increase was suppressed by BNIP3 knockdown. In addition, a similar increase in H3K27ac was observed with octanoate (Fig. 4j). These results suggest that BNIP3 promotes histone acetylation for p16 expression.

## Discussion

DNA damage has been proposed as the primary cause of aging, in part because most premature aging syndromes result from mutations in DNA repair genes^45^. Of note, these syndromes and normal aging are accompanied by mitochondrial dysfunction^45, 46^. DNA damage has been reported to affect mitochondrial metabolism and biogenesis by activating NAD^+^-consuming DNA repair enzymes and transcription factors of mitochondrial proteins^47^. In the present study, we uncovered a novel DDR signaling pathway that alters mitochondrial structure and metabolism to induce cellular senescence (Fig. 4k). Our results suggest that ATM activated by DNA damage translocates to mitochondria and phosphorylates BNIP3. BNIP3 remodels mitochondrial cristae, perhaps through interaction with MICOS, and activates FAO. This activation may promote histone acetylation for DNA repair^48^. However, if DNA damage persists, the resulting prolonged FAO activation appears to promote histone acetylation for p16 expression. The specificity to the *p16* gene is probably provided by transcription factors^32^. Although senescence-associated changes in mitochondrial metabolism have been previously reported, their significance is largely unknown^14^. Our results suggest that metabolic regulation of the epigenome is important for senescence induction. A recent study in DNA repair-deficient *C. elegans* showed that DNA damage shortens lifespan through FAO activation and histone hyperacetylation^49^. In addition, cristae morphology changes with age in mice and *Drosophila*^50^. We envision that DDR signaling to mitochondria via BNIP3 may be relevant to aging and age-related diseases.

## Methods

### Cell culture and treatments

IMR-90 normal human fibroblasts (ATCC, CCL-186) and HEK293T cells (ATCC, CRL-3216) were cultured in DMEM (Sigma–Aldrich, D5796; Wako, 044-29765) supplemented with 10% FBS (BioWest, S1560-500). Unless otherwise indicated, cells were treated with 200 ng ml^−1^ doxorubicin hydrochloride (Wako, 040-21521) or 5 mM sodium octanoate (Tokyo Chemical Industry, O0034, adjusted to pH 7.4 with HCl). MitoQ (MedChemExpress, HY-100116A) was added simultaneously with doxorubicin to a final concentration of 500 nM. To induce the expression of ER-KRAS^G12V^, 4-hydroxytamoxifen (Cayman Chemical, 17308) was added to a final concentration of 200 nM.

### Plasmids

To produce lentiviruses encoding shRNAs, shRNA sequences containing the following target sequences were cloned into the pLKO.1-TRC cloning vector (Addgene, 10878): nontargeting (NT), 5′-CAACAAGATGAAGAGCACCAA-3′; ATM #1, 5′-CCTGCCAACATACTTTAAGTA-3′; ATM #2, 5′-CGAGATCCTGAAACAATTAAA-3′; p53 #1, 5′-GACTCCAGTGGTAATCTAC-3′; p53 #2, 5′-GAGGGATGTTTGGGAGATGTA-3′; and BNIP3, 5′-GCTTCTGAAACAGATACCCAT-3′. To construct BNIP3 expression vectors, human BNIP3 was cloned into the pENTR or pENTR-5′-FLAG entry vector. Site-directed mutagenesis was performed to replace S2, S70, and S93 with alanine. To construct pENTR-shRNA-resistant BNIP3, site-directed mutagenesis was performed with the following primers: 5′-GATACCAACAGAGCAAGCGAAACAGATACCC-3′ and 5′-GGGTATCTGTTTCGCTTGCTCTGTTGGTATC-3′. LR reactions were performed using a Gateway LR Clonase enzyme mix (Invitrogen, 11791-100) and the pcDNA3-DEST or pLenti-PGK-Hygro (Addgene, 19066) destination vector. pLenti-PGK-ER-KRAS^G12V^ was purchased from Addgene (35635).

### Lentivirus infection

For lentivirus production, HEK293T cells were transfected with a lentiviral vector and the packaging plasmids pCMV-VSV-G (Addgene, 8454) and psPAX2 (Addgene, 12260) using PEI MAX transfection reagent (Polyscience, 24765) for 18 h. The medium was replaced with 5 ml of fresh medium containing 1% bovine serum albumin (BSA, Iwai Chemical, A001) and collected 24 h later. This step was repeated. The collected media were combined and filtered through a 0.45-μm pore size filter (Millipore, SLHVR33RS). IMR-90 cells were infected with lentiviruses in the presence of 8 μg ml^−1^ polybrene (Nacalai Tesque, 17736-44) for 18 h, cultured in fresh medium for 24 h, and selected with 0.5–1.0 μg ml^−1^ puromycin (Gibco, A11138-03) or 50–100 μg ml^−1^ hygromycin (Nacalai Tesque, 07296-11) for at least 2 days.

### Immunoblotting and immunoprecipitation

Cells were lysed in RIPA buffer (50 mM Tris-HCl [pH 8.0], 150 mM NaCl, 1% NP-40, 0.5% sodium deoxycholate, 0.1% sodium dodecyl sulfate [SDS], 1 mM phenylmethylsulfonyl fluoride, 5 µg ml^−1^ leupeptin, 8 mM NaF, 12 mM β-glycerophosphate, 1 mM Na_3_VO_4_, 1.2 mM Na_2_MoO_4_, 5 μM cantharidin, 2 mM imidazole) or IP lysis buffer (20 mM Tris-HCl [pH 7.5], 150 mM NaCl, 10 mM EDTA [pH 8.0], 1% Triton X-100, 1 mM phenylmethylsulfonyl fluoride, 5 µg ml^−1^ leupeptin, 8 mM NaF, 12 mM β-glycerophosphate, 1 mM Na_3_VO_4_, 1.2 mM Na_2_MoO_4_, 5 μM cantharidin, 2 mM imidazole). Mitochondria were isolated using a mitochondria isolation kit (Qiagen, 37612) according to the manufacturer’s protocol and lysed in RIPA buffer. The lysates were clarified by centrifugation at 13,000*g* for 15 min at 4 °C. The protein concentrations of the lysates were quantified using a BCA protein assay kit (Wako, 297-73101) and equalized. The lysates were then mixed with 2× SDS sample buffer (125 mM Tris-HCl [pH 6.8], 4% SDS, 20% glycerol, 200 μg ml^−1^ bromophenol blue, 10% β-mercaptoethanol) and heated at 98 °C for 3 min. Immunoprecipitation was performed with anti-FLAG M2 beads (Sigma–Aldrich, A2220). The beads were washed three times with IP lysis buffer and heated in 2× SDS sample buffer. The SDS samples were subjected to SDS polyacrylamide gel electrophoresis (SDS–PAGE) and transferred to Immobilon-P membranes (Millipore, IPVH00010). The membranes were blocked with 5% skim milk (Yukijirushi) in TBS-T (50 mM Tris-HCl [pH 8.0], 150 mM NaCl, 0.05% Tween 20) for 1 h at room temperature and incubated with primary antibodies in TBS-T containing 5% BSA and 0.1% NaN_3_ overnight at 4 °C. The membranes were then incubated with secondary antibodies in TBS-T containing 5% skim milk for 30 min at room temperature. ECL Select detection reagent (Amersham, RPN2235) was used for detection on a FUSION Solo S chemiluminescence imaging system (Vilber). The following primary antibodies were used: anti-p16 (Abcam, ab108349), anti-ATM (Abcam, ab32420), anti-α-tubulin (Bio-Rad, MCA77G), anti-phospho-ATM S1981 (Abcam, ab81292), anti-p21 (Abcam, ab109199), anti-p53 (Santa Cruz Biotechnology, sc-126), anti-lamin B1 (Abcam, ab16048), anti-BNIP3 (Cell Signaling Technology, 44060), anti-GAPDH (Proteintech, 60004-1-Ig), anti-NDUFA8 (Abcam, ab184952), anti-β-actin (Sigma-Aldrich, A3853), anti-KRAS (Santa Cruz Biotechnology, sc-30), anti-cytochrome *c* (BD Biosciences, 556433), anti-histone H3 (Abcam, ab1791), anti-phospho-ATM/ATR substrate (SQ) (Cell Signaling Technology, 9607), anti-MIC60 (Proteintech, 10179-1-AP), anti-MIC19 (Proteintech, 25625-1-AP), anti-PDH E1α (Santa Cruz Biotechnology, sc-377092), anti-phospho-PDH E1α S293 (Abcam, ab177461), and anti-CPT2 (Santa Cruz Biotechnology, sc-377294). A phospho-specific antibody against BNIP3 S70 was raised by immunizing a rabbit with a keyhole limpet hemocyanin-conjugated phosphopeptide (CDSPPR[pS]QTPQD) and affinity-purified (Eurofins).

### qPCR

Total RNA was extracted using Isogen (Wako, 319-90211) and subjected to reverse transcription using a ReverTra Ace master mix (Toyobo, FSQ-301). qPCR was performed using a SYBR FAST qPCR kit (KAPA Biosystems, KK4602) and a QuantStudio 1 qPCR system (Applied Biosystems). The following primers were used: *GAPDH*, forward, 5′-AGCCACATCGCTCAGACAC-3′, reverse, 5′-GCCCAATACGACCAAATCC-3′; *p16*, forward, 5′-CCAACGCACCGAATAGTTACG-3′, reverse, 5′-GCGCTGCCCATCATCATG-3′; *BNIP3*, forward, 5′-TGCTGCTCTCTCATTTGCTG-3′, reverse, 5′-GACTCCAGTTCTTCATCAAAAGGT-3′; *NDUFA8*, forward, 5′-ATGCCGGGGATAGTGGAG-3′, reverse, 5′-GCACAGCAGAACTAATTTTCACC-3′; *IL1A*, forward, 5′-GCCAGCCAGAGAGGGAGTC-3′, reverse, 5′-TGGAACTTTGGCCATCTTGAC-3′; *IL1Β*, forward, 5′-TACCTGTCCTGCGTGTTGAA-3′, reverse, 5′-TCTTTGGGTAATTTTTGGGATCT-3′; *IL6*, forward, 5′-CCGGGAACGAAAGAGAAGCT-3′, reverse, 5′-GCGCTTGTGGAGAAGGAGTT-3′; *CCL2*, forward, 5′-AGTCTCTGCCGCCCTTCT-3′, reverse, 5′-GTGACTGGGGCATTGATTG-3′; and *IL8*, forward, 5′-CTTTCCACCCCAAATTTATCAAAG-3′, reverse, 5′-CAGACAGAGCTCTCTTCCATCAGA-3′. Transcript levels were analyzed using the 1′1′*C*_t_ method with *GAPDH* as the internal control.

### siRNA transfection

siRNAs with the following target sequences were purchased from Dharmacon: NT (siGENOME), 5′-AUGAACGUGAAUUGCUCAA-3′; ATM (siGENOME), 5′-GCAAAGCCCUAGUAACAUA-3′; p16 (siGENOME), 5′-AAACUUAGAUCAUCAGUCA-3′; NT #1 (hereafter ON-TARGETplus), 5′-UGGUUUACAUGUCGACUAA-3′; NT #2, 5′-UGGUUUACAUGUUGUGUGA-3′; BNIP3 #1, 5′-GGAAAGAAGUUGAAAGCAU-3′; BNIP3 #2, 5′-GGAAUUAAGUCUCCGAUUA-3′; NDUFA8 #1, 5′-GAAAACAGAUCGACCUUUA-3′; NDUFA8 #2, 5′-GUAGAUGAGGUGAAAAUUA-3′; ATM, 5′-GCAAAGCCCUAGUAACAUA-3′; MIC60 #1, 5′-GGGAUGACUUUAAACGAGA-3′; MIC60 #2, 5′-CAGACAAACUCUUCGAGAU-3′; CPT2 #1, 5′-CUAGAUGACUUCCCCAUUA-3′; CPT2 #2, 5′-GGGCCUACCUGGUCAAUGC-3′; and p16, 5′-GAUCAUCAGUCACCGAAGG-3′. ON-TARGETplus and the #1 siRNAs were used unless otherwise indicated. siRNA transfection was performed at a final concentration of 10 nM using Lipofectamine RNAiMAX transfection reagent (Invitrogen, 13778500) and Opti-MEM (Gibco, 31985070) for 24–48 h.

### Genome-wide siRNA screening

siGENOME SMARTpool siRNA libraries (Human Drug Targets, G-004655-E2; Human Druggable Subsets, G-004675-E2; Human Genome, G-005005-E2) were purchased from Dharmacon. The siRNA in each well was diluted to 375 nM in siRNA buffer (Dharmacon, B-002000-UB-100). A total of 1.5 pmol of siRNA (4 μl) was dispensed into each well of black clear-bottom 384-well plates (Greiner, 781091) with a Biomek FX^P^ liquid handler (Beckman Coulter). Lipofectamine RNAiMAX transfection reagent and Opti-MEM were mixed at a volume ratio of 3:400 and added to each well using a Multidrop Combi automated pipetting machine (Thermo Scientific). IMR-90 cells (population doubling level 40 or lower, 1,600 cells per well) were transfected with siRNAs at a final concentration of 30 nM for 24 h and treated with 250 ng ml^−1^ doxorubicin for 10 days. For immunofluorescence, cells were fixed with 4% paraformaldehyde in PBS for 10 min and permeabilized with 0.1% Triton X-100 in PBS containing 1% BSA for 5 min. The cells were washed with PBS using an AquaMax microplate washer (Molecular Devices) and incubated with an anti-p16 antibody (1:1000, BD Biosciences, 551154) in PBS containing 1% BSA overnight at 4 °C. After washing with PBS, the cells were incubated with Alexa Fluor Plus 647 (1:1000, Invitrogen, A32728) and Hoechst 33258 (1:2000, Dojindo, H341) in PBS containing 1% BSA for 1 h at room temperature and washed with PBS. Images were acquired with a CellInsight NXT automated microscope (Thermo Scientific) and analyzed with HCS Studio software (Thermo Scientific). The mean fluorescence intensity of p16 in the nucleus was quantified and log-transformed to achieve a near-normal distribution. The robust Z-score of each siRNA was calculated. siRNAs with cell counts below 30 were excluded from the analysis.

### EdU assay

A Click-iT EdU imaging kit (Invitrogen, C10340) was used. IMR-90 cells were incubated with 10 µM EdU for 24 h, fixed with 4% paraformaldehyde in PBS for 15 min at room temperature, and washed twice with PBS containing 3% BSA. The cells were permeabilized with 0.5% Triton X-100 in PBS for 20 min at room temperature and washed with PBS containing 3% BSA. Click-iT reaction cocktail was added and allowed to react for 30 min at room temperature. After washing with PBS containing 3% BSA, the cells were incubated with Hoechst 33342 (1:2000) in PBS for 30 min at room temperature. Images were acquired with a CellInsight NXT automated microscope and analyzed with HCS Studio software.

### Measurement of ROS

IMR-90 cells were incubated with 5 μM MitoSOX mitochondrial superoxide indicator (Invitrogen, M36008) and Hoechst 33342 (1:1000, Dojindo, H342) in serum-free DMEM or with 2.5 μM CellROX ROS indicator (Invitrogen, C10422) and Hoechst 33342 (1:1000) in culture medium for 30 min at 37 °C and washed with PBS. To evaluate and subtract autofluorescence, cells were also stained with Hoechst 33342 only. Images were acquired with a CellInsight NXT automated microscope and analyzed with HCS Studio software.

### Immunofluorescence

IMR-90 cells were grown on glass coverslips (Matsunami Glass, C015001), fixed with 4% paraformaldehyde in PBS for 10 min, and permeabilized with 0.1% Triton X-100 in PBS containing 1% BSA for 5 min. The cells were incubated with anti-BNIP3 (Abcam, ab109362) and anti-TOM20 (Santa Cruz Biotechnology, sc-17764) antibodies in PBS containing 1% BSA for 1 h and then with Alexa Fluor secondary antibodies (1:1000, Invitrogen) and Hoechst 33258 (1:2000) for 1 h. Images were acquired using a TCS SP5 confocal microscope (Leica) with a 63× oil immersion objective (NA 1.40, Leica).

### Protein mass spectrometry

HEK293T cells were transfected with pcDNA3 empty vector and pcDNA3-FLAG-BNIP3 in 10 cm dishes for 24 h, lysed in IP lysis buffer, and centrifuged at 15,000*g* for 15 min at 4 °C. The supernatants were incubated with FLAG M2 beads for 30 min at 4 °C. The beads were washed with IP lysis buffer and incubated with 5 mg ml^−1^ 3× FLAG peptide (Sigma–Aldrich, F4799; M&S TechnoSystems, GEN-3-FLAG-5) in TBS (10 mM Tris-HCl [pH 7.4], 150 mM NaCl) for 30 min at 4 °C. After centrifugation, the supernatants were collected in low-protein-binding tubes (Eppendorf, 0030108442). Trichloroacetic acid was added to the supernatants to a final concentration of 10%. The supernatants were incubated on ice for 30 min and centrifuged at 15,000*g* for 10 min at 4 °C. The precipitates were resuspended in 200 μl of ice-cold acetone and centrifuged at 15,000*g* for 10 min at 4 °C. The precipitates were then resuspended in ultrapure water (Wako, 214-01301) containing 50 mM ammonium bicarbonate and 0.1% RapiGest surfactant (Waters, 186002123) and heated for 10 min at 98 °C. The samples were shaken for 5 min and subjected to ultrasonication. The samples were then treated with 5 mM Tris(2-carboxyethyl)phosphine for 10 min at 98 °C, 10 mM methyl methanethiosulfonate for 30 min at room temperature, and 1 μg of mass spectrometry-grade trypsin (Promega, V5280) overnight at 37 °C. Trifluoroacetic acid was added to reach 0.5%. The samples were incubated for 40 min at 37 °C and centrifuged at 20,000*g* for 10 min at 4 °C. The supernatants were desalted using a GL-Tip SDB kit (GL Sciences, 7820-11200) according to the manufacturer’s protocol, except that the columns were washed twice with 200 μl of solution A. For phosphorylation analysis, phosphopeptides were enriched using a Titansphere Phos-TiO kit (GL Sciences, 5010-21305) prior to desalting. After vacuum centrifugation, the peptides were dissolved in 0.1% formic acid for protein identification or in 0.1% formic acid and 2% acetonitrile for phosphorylation analysis, clarified by centrifugation at 20,000*g* for 10 min at room temperature, and transferred to ultra-low-adsorption vials (AMR, PSVial100).

LC–MS/MS analysis was performed on a Q Exactive mass spectrometer (Thermo Scientific) connected to an EASY-nLC 1200 system (Thermo Scientific). Peptides were loaded onto a C_18_ trap column (Thermo Scientific, 164946) and a C_18_ analytical column (Nikkyo Technos, NTCC-360/75-3-123) and separated at a flow rate of 300 nl min^−1^ with a gradient of mobile phase A (0.1% formic acid in ultrapure water) and mobile phase B (0.1% formic acid in acetonitrile) according to the following protocols: 0–10% B for 5 min, 10–40% B for 85 min, 40–95% B for 2 min, and 95% B for 18 min for protein identification; and 2–40% B for 85 min, 40–95% B for 5 min, 95% B for 10 min, 95–2% B for 2 min, and 2% B for 8 min for phosphorylation analysis. The following MS settings were used: range, 400–1,500 *m/z* (protein identification) or 350–1,500 *m/z* (phosphorylation analysis); resolution, 70,000; automatic gain control target, 3E6; and maximum injection time, 60 ms. The following MS^2^ settings were used: topN, 10; normalized collision energy, 28 (protein identification) or 27 (phosphorylation analysis); isolation window, 3.0 *m/z* (protein identification) or 1.6 *m/z* (phosphorylation analysis); resolution, 17,500; automatic gain control target, 1E5 (protein identification) or 5E5 (phosphorylation analysis); and maximum injection time, 60 ms (protein identification) or 100 ms (phosphorylation analysis). The data were analyzed with Proteome Discoverer software (Thermo Scientific). GO analysis was performed using DAVID Bioinformatics Resources.

### Electron microscopy

IMR-90 cells were fixed with 2% paraformaldehyde and 2% glutaraldehyde in 0.1 M phosphate buffer (pH 7.4) overnight at 4 °C and postfixed with 2% osmium tetroxide for 2 h at 4 °C. The cells were washed in distilled water for 10 min three times and block-stained with 1% uranyl acetate for 30 min. The cells were then washed in distilled water three times, dehydrated with a graded series of ethanol, and embedded in Epon 812 resin (Oken Shoji) for 72 h at 60 °C. Ultrathin sections (100 nm) were cut with a Leica EM UC6 ultramicrotome (Leica), placed on 12 mm circular glass coverslips, and stained with uranyl acetate and lead citrate. The coverslips were mounted on aluminum pin stubs with carbon tape (Nissin EM) and carbon paste (Pelco Colloidal Graphite) and coated using a carbon coater (CADE-4T, Meiwafosis) to prevent electron charging. Electron micrographs were acquired using a Helios Nanolab 660 FIB–SEM (Thermo Scientific) equipped with a backscattered electron detector (circular backscatter detector) at an acceleration voltage of 2.0 kV and a current of 0.4 nA.

### FIB–SEM

Resin-embedded samples were glued to aluminum pin stubs, cut with a Leica EM UC6 ultramicrotome, and coated with a thin layer of Pt–Pd using an MC1000 ion sputter coater (Hitachi High Technologies). Serial electron micrographs were acquired every 10 nm using a Helios Nanolab 660 FIB–SEM equipped with a backscattered electron detector (MD detector) at an acceleration voltage of 2.0 kV and a current of 0.4 nA. FIB milling was performed at an acceleration voltage of 30 kV with a beam current of 0.4 nA. 3D reconstruction was performed using Amira software (Thermo Scientific).

### Metabolomic analysis

IMR-90 cells were grown in 10 cm dishes, washed twice with ice-cold PBS, and lysed in 1 ml of methanol containing 25 µM L-methionine sulfone and 25 µM MES as internal standards and 400 µL of ultrapure water. The lysate (1 ml) was mixed with 800 µL of chloroform and centrifuged at 10,000*g* for 3 min at 4 °C. The aqueous layer was transferred to a centrifugal filter (Human Metabolome Technologies, UFC3LCCNB-HMT) and centrifuged at 9,100*g* for 3 h at 4 °C. The solvent of the filtrate was evaporated by blowing nitrogen gas for 2 h at 40 °C (TAITEC). The residue was dissolved in 25 µl of ultrapure water. The levels of anionic metabolites were quantified using a Q Exactive Focus mass spectrometer (Thermo Scientific) connected to an ICS-5000+ chromatography compartment (Thermo Scientific). The samples were separated on a Dionex IonPac AS11-HC column (4 μm particle size, Thermo Scientific) at a flow rate of 0.25 ml min^−1^ with a gradient of potassium hydroxide according to the following protocol: 1 mM to 100 mM (0–40 min), 100 mM (40–50 min), and 1 mM (50–60 min). The column temperature was set at 30 °C. A post-column make-up flow of methanol was added at a flow rate of 0.18 ml min^−1^. The potassium hydroxide gradient was replaced with pure water before entering the mass spectrometer using a Dionex AERS 500 anion electrolytic suppressor (Thermo Scientific). The mass spectrometer was operated in ESI negative mode. A full mass scan was performed with the following settings: range, 70–900 *m/z*; resolution, 70,000; automatic gain control target, 3E6; and maximum ion injection time, 100 ms. The ion source parameters were as follows: spray voltage, 3 kV; transfer temperature, 320 °C; S-lens level, 50; heater temperature, 300 °C; sheath gas, 36; and aux gas, 10. The levels of cationic metabolites were quantified by LC–MS/MS. The samples were separated on a Discovery HS F5-3 column (2.1 mm i.d. × 150 mm long, 3 µm particle size, Sigma–Aldrich) at a flow rate of 0.25 ml min^−1^ with a step gradient of mobile phase A (0.1% formic acid) and mobile phase B (0.1% acetonitrile) according to the following protocol: 100:0 (0–5 min), 75:25 (5–11 min), 65:35 (11–15 min), 5:95 (15–20 min), and 100:0 (20–25 min).

The column temperature was set at 40 °C. An LCMS-8060 triple quadrupole mass spectrometer (Shimadzu) equipped with an ESI ion source was operated in positive and negative ESI, multiple reaction monitoring (MRM) mode. The relative level of each metabolite was normalized to the protein concentration of duplicate samples as measured by BCA assay. For isotope labeling, DMEM without glucose and glutamine (Sigma–Aldrich, D5030) was supplemented with [U-^13^C]glucose (Sigma–Aldrich, 389374) and glutamine (Sigma–Aldrich, G-9723).

### Measurement of FAO activity

IMR-90 cells were seeded in black glass-bottom 96-well plates (Matsunami Glass, GP96000), washed twice with Hanks’ balanced solution (HBS; 20 mM HEPES, 107 mM NaCl, 6 mM KCl, 1.2 mM MgSO_4_, 2 mM CaCl_2_, 11.5 mM glucose, pH 7.4) and incubated with 20 μM FAOBlue (Funakoshi, FDV-0033) in HBS for 2 h at 37 °C. To evaluate and subtract autofluorescence, cells were also incubated in HBS without FAOBlue. Fluorescence was measured with a Varioskan microplate reader (Ex/Em = 405/460 nm, Thermo Scientific). To normalize measurements to cell number, cells were fixed with methanol for 5 min, stained with Hoechst 33258 (1:2000) in PBS for 30 min, and counted with a CellInsight NXT automated microscope.

### Lipid droplet staining

IMR-90 cells were seeded on glass coverslips, fixed with 4% paraformaldehyde in PBS for 10 min, and incubated with 1 μM BODIPY 493/503 (Cayman Chemical, 25892) and DAPI (1:2000, Dojindo, D523). Images were acquired using a TCS SP5 confocal microscope with a 63× oil immersion objective. Lipid droplet size was quantified using the ‘‘analyze particles’’ tool of ImageJ software (NIH).

### SA-β-gal staining

An SA-β-gal staining kit (Cell Signaling Technology, 9860) was used according to the manufacturer’s protocol.

### ChIP–qPCR

ChIP was performed using anti-H3K27ac (Active Motif, 39133) and anti-histone H3 (Abcam, ab1791) antibodies and an enzymatic ChIP kit (Cell Signaling Technology, 9003) according to the manufacturer’s protocol. qPCR was performed with the following primers: primer pair 1, forward, 5′-GGATTCTAAGCCAACATCATTTC-3′, reverse, 5′-TGGATAGTTTTGACAATTTTTAATGG-3′; and primer pair 2, forward, 5′-TCCCCTCTGATATTATTAAAGATTGC-3′, reverse, 5′-GTTGAATCACATATCAGGTGAAGAAT-3′. The percent input was calculated using the 1′1′*C*t method. ChIP-seq data (GSM1915113, GSM1915114, GSM1915115) were downloaded from the Gene Expression Omnibus database and visualized with Integrative Genomics Viewer (Broad Institute).

### Statistical analyses

R software and Microsoft Excel were used for statistical analyses. Statistical tests and sample size are reported in the figure legends. *P* < 0.05 was considered statistically significant. No statistical method was used to predetermine the sample size.

## Acknowledgements

We thank A. Takahashi for reagents and advice on senescence experiments; T. Fujisawa for assistance with analysis of ChIP–seq data; and S. Hirayama for assistance with mass spectrometry. This work was supported by grants from the Japan Agency for Medical Research and Development (JP21gm5010001 to H.I.), the Japan Science and Technology Agency (JPMJMS2022-18 to H.I.), and the Japan Society for the Promotion of Science (JP18H03995 and JP21H04760 to H.I. and 18K14648 and 21K06061 to S.Y.).

## Author contributions

S.Y. conceived the study, designed the experiments, performed most of the experiments, analyzed the data, and wrote the manuscript. Y.S. performed metabolomic analysis. J.Y. and Y.U. performed electron microscopy. X.Z., T.O., and S.F. performed experiments. I.N. helped perform experiments. H.I. conceived the study and wrote the manuscript.

## Competing interests

The authors declare no competing interests.

**Correspondence and requests for materials** should be addressed to S.Y. or H.I.

---

**Extended Data Fig. 1.**
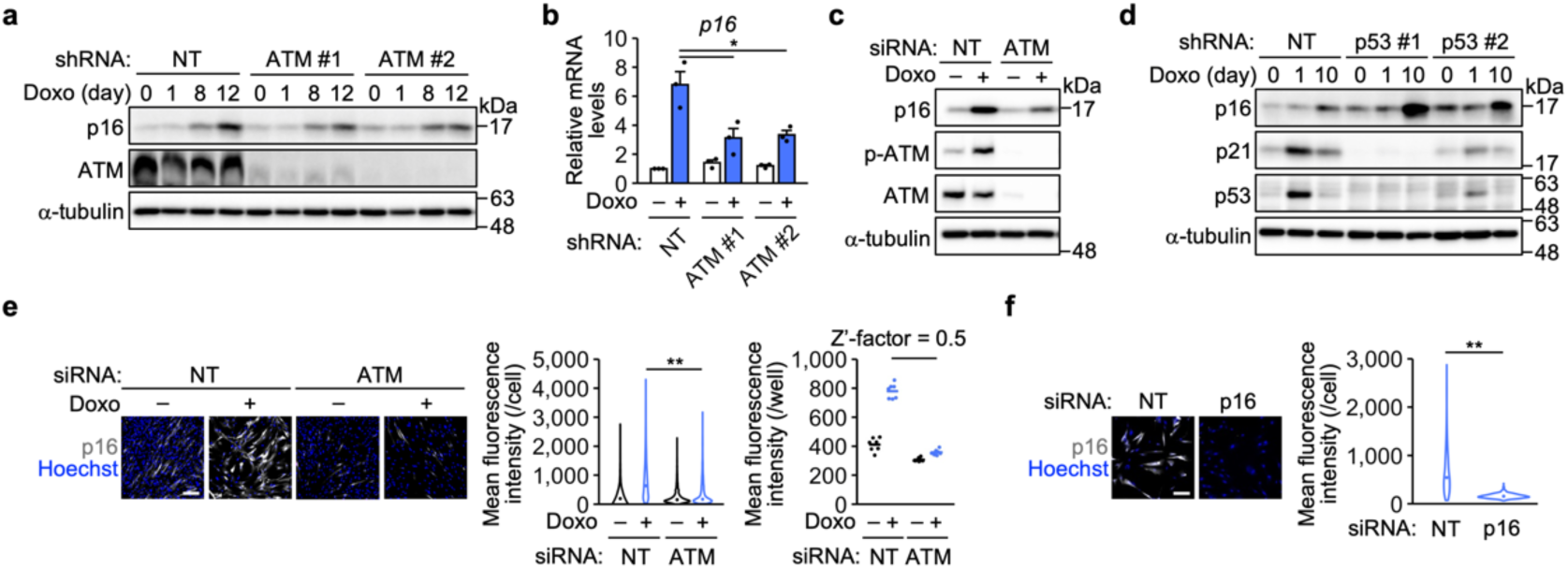
p16 expression requires ATM but not p53. **a**, Immunoblot analysis of IMR-90 cells infected with lentiviruses encoding nontargeting (NT) and ATM shRNAs and treated with doxorubicin (Doxo). Representative of two independent experiments. **b**, qPCR analysis of IMR-90 cells infected with lentiviruses encoding NT and ATM shRNAs and treated with Doxo for 12 days. The bars represent the means ± s.e.m. *n* = 3 independent experiments. **c**, Immunoblot analysis of IMR-90 cells transfected with NT and ATM siRNAs and treated with Doxo for 12 days. Representative of three independent experiments. **d**, Immunoblot analysis of IMR-90 cells infected with lentiviruses encoding NT and p53 shRNAs and treated with Doxo. Representative of three independent experiments. **e**, Left, immunofluorescence analysis of IMR-90 cells transfected with NT and ATM siRNAs (siGENOME) and treated with Doxo for 10 days. Center, distribution of the mean fluorescence intensity per cell. The dots represent the median values. *n* = 3405 (siNT), 1099 (siNT + Doxo), 2926 (siATM), 1306 (siATM + Doxo) cells. Right, mean fluorescence intensity per well. The lines represent the mean values. *n* = 7 wells. **f**, Left, immunofluorescence analysis of Doxo-induced senescent cells transfected with NT and p16 siRNAs (siGENOME). Right, distribution of the mean fluorescence intensity per cell. The dots represent the median values. *n* = 704 (siNT), 755 (sip16) cells. Scale bars, 200 μm. Statistical analysis was performed using Dunnett’s multiple comparison test (**b**) and the Wilcoxon rank-sum test with Bonferroni correction (**e**, center, **f**). \**P* < 0.05, ** *P* < 0.01.

**Extended Data Fig. 2.**
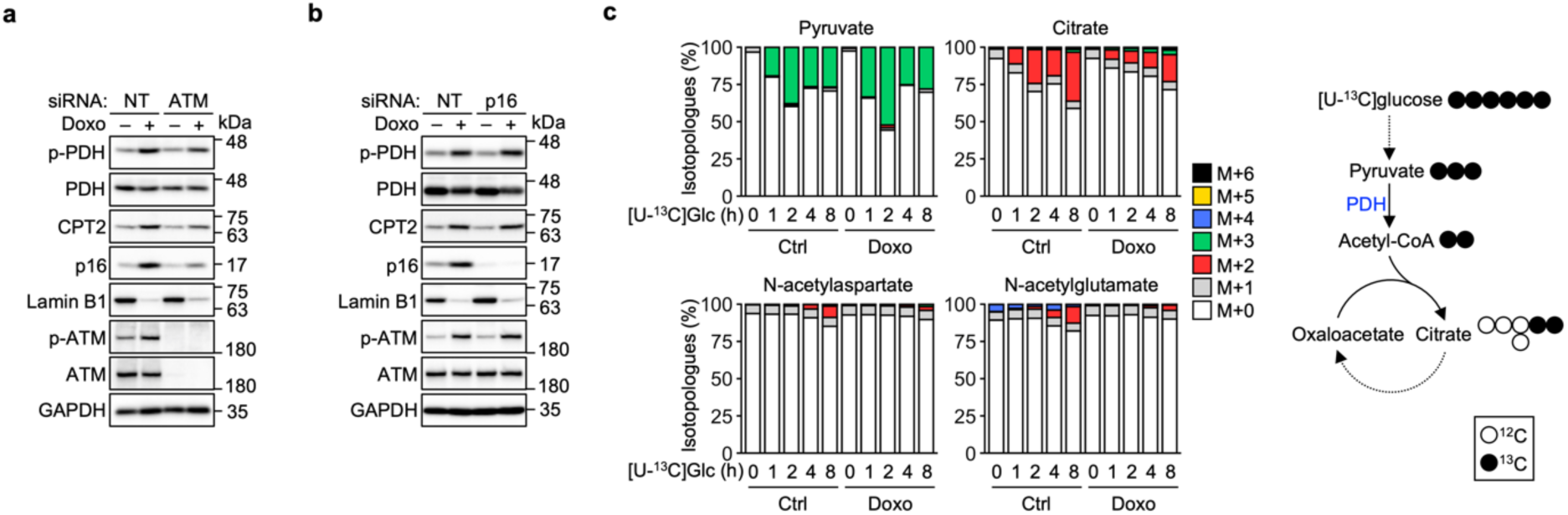
PDH activity decreases during DNA damage-induced senescence. **a**, Immunoblot analysis of IMR-90 cells transfected with NT and ATM siRNAs and treated with doxorubicin (Doxo) for 12 days. Representative of two independent experiments. **b**, Immunoblot analysis of IMR-90 cells transfected with NT and p16 siRNAs and treated with Doxo for 12 days. Representative of two independent experiments. **c,** Left, isotope labeling in control and Doxo-induced senescent cells. *n* = 1 biological replicate for each time point. Right, isotope labeling patterns associated with PDH activity.

**Extended Data Fig. 3.**
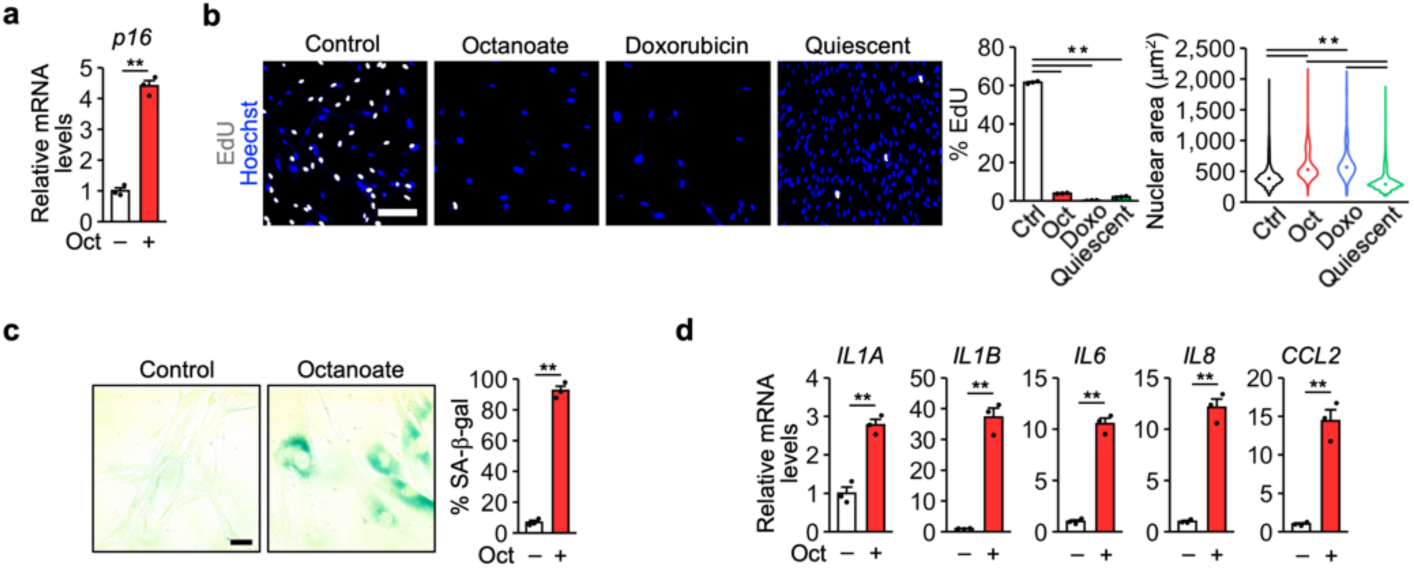
Octanoate induces senescence-like proliferation arrest. **a**, **d**, qPCR analysis of IMR-90 cells treated with octanoate (Oct) for 12 days. *n* = 3 biological replicates. **b**, Left, EdU assay using IMR-90 cells treated with Oct and doxorubicin (Doxo) for 12 days or made quiescent by confluence. Scale bar, 200 μm. Center, percentage of EdU-positive cells. *n* = 3 biological replicates. Right, distribution of the nuclear area. The dots represent the median values. *n* = 7790 (Ctrl), 3748 (Oct), 967 (Doxo), 17185 (quiescent) cells. **c**, SA-β-gal staining in IMR-90 cells treated with Oct for 12 days. Scale bar, 50 μm. *n* = 3 biological replicates. Statistical analysis was performed using unpaired two-tailed Student’s *t*-test (**a**, **c**, **d**), Dunnett’s multiple comparison test (**b**, center), and the Wilcoxon rank-sum test with Bonferroni correction (**b**, right). ** *P* < 0.01.

## References

1. Di Micco, R., Krizhanovsky, V., Baker, D. & d’Adda di Fagagna, F. Cellular senescence in ageing: from mechanisms to therapeutic opportunities. Nat. Rev. Mol. Cell Biol. 22, 75–95 (2021).

2. Baker, D. J. et al. Naturally occurring p16^Ink4a^-positive cells shorten healthy lifespan. Nature 530, 184–189 (2016).

3. Serrano, M., Lin, A. W., McCurrach, M. E., Beach, D. & Lowe, S. W. Oncogenic *ras* provokes premature cell senescence associated with accumulation of p53 and p16^INK4a^. Cell 88, 593–602 (1997).

4. Yamakoshi, K. et al. Real-time in vivo imaging of p16^Ink4a^ reveals cross talk with p53. J. Cell Biol. 186, 393–407 (2009).

5. Burd, C. E. et al. Monitoring tumorigenesis and senescence in vivo with a *p16^INK4a^*-luciferase model. Cell 152, 340–351 (2013).

6. Krishnamurthy, J. et al. p16^INK4a^ induces an age-dependent decline in islet regenerative potential. Nature 443, 453–457 (2006).

7. Molofsky, A. V. et al. Increasing *p16^INK4a^*expression decreases forebrain progenitors and neurogenesis during ageing. Nature 443, 448–452 (2006).

8. Sharpless, N. E. & Sherr, C. J. Forging a signature of *in vivo* senescence. Nat. Rev. Cancer 15, 397–408 (2015).

9. LaPak, K. M. & Burd, C. E. The molecular balancing act of p16^INK4a^ in cancer and aging. Mol. Cancer Res. 12, 167–183 (2014).

10. Kang, C. et al. The DNA damage response induces inflammation and senescence by inhibiting autophagy of GATA4. Science 349, aaa5612 (2015).

11. Zhang, J., Chung, T. & Oldenburg, K. A simple statistical parameter for use in evaluation and validation of high throughput screening assays. J. Biomol. Screen. 4, 67–73 (1999).

12. Zhang, J. & Ney, P. A. Role of BNIP3 and NIX in cell death, autophagy, and mitophagy. Cell Death Differ. 16, 939–946 (2009).

13. Stroud, D. A. et al. Accessory subunits are integral for assembly and function of human mitochondrial complex I. Nature 538, 123–126 (2016).

14. Martini, H. & Passos, J. F. Cellular senescence: all roads lead to mitochondria. FEBS J. in press

15. Correia-Melo, C. et al. Mitochondria are required for pro-ageing features of the senescent phenotype. EMBO J. 35, 724–742 (2016).

16. Xu, D. & Finkel, T. A role for mitochondria as potential regulators of cellular life span. Biochem. Biophys. Res. Commun. 294, 245–248 (2002).

17. Wiley, C. D. et al. Mitochondrial dysfunction induces senescence with a distinct secretory phenotype. Cell Metab. 23, 303–314 (2016).

18. Moiseeva, O., Bourdeau, V., Roux, A., Deschenes-Simard, X. & Ferbeyre, G. Mitochondrial dysfunction contributes to oncogene-induced senescence. Mol. Cell. Biol. 29, 4495–4507 (2009).

19. Nacarelli, T. et al. NAD^+^ metabolism governs the proinflammatory senescence-associated secretome. Nat. Cell Biol. 21, 397–407 (2019).

20. Vizioli, M. G. et al. Mitochondria-to-nucleus retrograde signaling drives formation of cytoplasmic chromatin and inflammation in senescence. Genes Dev. 34, 428–445 (2020).

21. Chourasia, A. H. et al. Mitophagy defects arising from BNip3 loss promote mammary tumor progression to metastasis. EMBO Rep. 16, 1145–1163 (2015).

22. Dou, Z. et al. Cytoplasmic chromatin triggers inflammation in senescence and cancer. Nature 550, 402–406 (2017).

23. Takahashi, A. et al. Downregulation of cytoplasmic DNases is implicated in cytoplasmic DNA accumulation and SASP in senescent cells. Nat. Commun. 9, 1249 (2018).

24. Georgilis, A. et al. PTBP1-mediated alternative splicing regulates the inflammatory secretome and the pro-tumorigenic effects of senescent cells. Cancer Cell 34, 85–102.e9 (2018).

25. Yamauchi, S. et al. p53-mediated activation of the mitochondrial protease HtrA2/Omi prevents cell invasion. J. Cell Biol. 204, 1191–1207 (2014).

26. Shiloh, Y. & Ziv, Y. The ATM protein kinase: regulating the cellular response to genotoxic stress, and more. Nat. Rev. Mol. Cell Biol. 14, 197–210 (2013).

27. Valentin-Vega, Y. A. et al. Mitochondrial dysfunction in ataxia-telangiectasia. Blood 119, 1490–1500 (2012).

28. Mai, S., Klinkenberg, M., Auburger, G., Bereiter-Hahn, J. & Jendrach, M. Decreased expression of Drp1 and Fis1 mediates mitochondrial elongation in senescent cells and enhances resistance to oxidative stress through PINK1. J. Cell Sci. 123, 917–926 (2010).

29. Gomes, L. C., Di Benedetto, G. & Scorrano, L. During autophagy mitochondria elongate, are spared from degradation and sustain cell viability. Nat. Cell Biol. 13, 589–598 (2011).

30. Rampelt, H., Zerbes, R. M., van der Laan, M. & Pfanner, N. Role of the mitochondrial contact site and cristae organizing system in membrane architecture and dynamics. Biochim. Biophys. Acta 1864, 737–746 (2017).

31. Gok, M. O. & Friedman J. R. The outer mitochondrial membrane protein TMEM11 is a novel negative regulator of BNIP3/BNIP3L-dependent receptor-mediated mitophagy. Preprint at https://www.biorxiv.org/content/10.1101/2022.03.29.486240v1 (2022).

32. Martínez-Reyes, I. & Chandel, N. S. Mitochondrial TCA cycle metabolites control physiology and disease. Nat. Commun. 11, 102 (2020).

33. Bartlett, K. & Eaton, S. Mitochondrial β-oxidation. Eur. J. Biochem. 271, 462– 469 (2004).

34. Buck, M. D. et al. Mitochondrial dynamics controls T cell fate through metabolic programming. Cell 166, 63–76 (2016).

35. Chung, S. et al. Mitochondrial oxidative metabolism is required for the cardiac differentiation of stem cells. Nat. Clin. Pract. Cardiovasc. Med. 4 **Suppl 1**, S60–7 (2007).

36. Rambold, A. S., Cohen, S. & Lippincott-Schwartz, J. Fatty acid trafficking in starved cells: regulation by lipid droplet lipolysis, autophagy, and mitochondrial fusion dynamics. Dev. Cell 32, 678–692 (2015).

37. Alan, L. & Scorrano, L. Shaping fuel utilization by mitochondria. Curr. Biol. 32, R618–R623 (2022).

38. Quijano, C. et al. Oncogene-induced senescence results in marked metabolic and bioenergetic alterations. Cell Cycle 11, 1383–1392 (2012).

39. Glick, D. et al. BNip3 regulates mitochondrial function and lipid metabolism in the liver. Mol. Cell. Biol. 32, 2570–2584 (2012).

40. Hue, L. & Taegtmeyer, H. The Randle cycle revisited: a new head for an old hat. Am. J. Physiol. Endocrinol. Metab. 297, E578–91 (2009).

41. McDonnell, E. et al. Lipids reprogram metabolism to become a major carbon source for histone acetylation. Cell Rep. 17, 1463–1472 (2016).

42. Munro, J., Barr, N. I., Ireland, H., Morrison, V. & Parkinson, E. K. Histone deacetylase inhibitors induce a senescence-like state in human cells by a p16-dependent mechanism that is independent of a mitotic clock. Exp. Cell Res. 295, 525–538 (2004).

43. Tasdemir, N. et al. BRD4 connects enhancer remodeling to senescence immune surveillance. Cancer Discov. 6, 612–629 (2016).

44. Haberle, V. & Stark, A. Eukaryotic core promoters and the functional basis of transcription initiation. Nat. Rev. Mol. Cell Biol.19, 621–637 (2018).

45. Yousefzadeh, M. et al. DNA damage-how and why we age? eLife 10, e62852 (2021).

46. Wang, Y. & Hekimi, S. Mitochondrial dysfunction and longevity in animals: Untangling the knot. Science 350, 1204–1207 (2015).

47. Fang, E. F. et al. Nuclear DNA damage signalling to mitochondria in ageing. Nat. Rev. Mol. Cell Biol. 17, 308–321 (2016).

48. Sivanand, S. et al. Nuclear Acetyl-CoA production by ACLY promotes homologous recombination. Mol. Cell 67, 252–265.e6 (2017).

49. Hamsanathan, S. et al. Integrated -omics approach reveals persistent DNA damage rewires lipid metabolism and histone hyperacetylation via MYS-1/Tip60. Sci. Adv. 8, eabl6083 (2022).

50. Brandt, T. et al. Changes of mitochondrial ultrastructure and function during ageing in mice and Drosophila. eLife 6, e24662 (2017).

